# RAREFAN: A webservice to identify REPINs and RAYTs in bacterial genomes

**DOI:** 10.1101/2022.05.22.493013

**Authors:** Carsten Fortmann-Grote, Julia von Irmer, Frederic Bertels

**Author notes:** Corresponding author: Frederic Bertels, August-Thienemann-Straße 2, 24306 Plön, Germany,.

## Abstract

Compared to eukaryotes, repetitive sequences are rare in bacterial genomes and usually do not persist for long. Yet, there is at least one class of persistent prokaryotic mobile genetic elements: REPINs. REPINs are non-autonomous transposable elements replicated by single-copy transposases called RAYTs. REPIN-RAYT systems are mostly vertically inherited and have persisted in individual bacterial lineages for millions of years. Discovering and analyzing REPIN populations and their corresponding RAYT transposases in bacterial species can be rather laborious, hampering progress in understanding REPIN-RAYT biology and evolution. Here we present RAREFAN, a webservice that identifies REPIN populations and their corresponding RAYT transposase in a given set of bacterial genomes. We demonstrate RAREFAN’s capabilities by analyzing a set of 49 *Stenotrophomonas maltophilia* genomes, containing nine different REPIN-RAYT systems. We guide the reader through the process of identifying and analyzing REPIN-RAYT systems across *S. maltophilia*, highlighting erroneous associations between REPIN and RAYTs, and providing solutions on how to find correct associations. RAREFAN enables rapid, large-scale detection of REPINs and RAYTs, and provides insight into the fascinating world of intragenomic sequence populations in bacterial genomes. RAREFAN is available at http://rarefan.evolbio.mpg.de.

## Introduction

Repetitive sequences in bacteria are rare compared to most eukaryotic genomes. In eukaryotic genomes, repetitive sequences are the result of the activities of persistent parasitic transposable elements. In bacteria, in contrast, parasitic transposable elements cannot persist for long periods of time (Park *et al*. 2021; van Dijk *et al*. 2022). To persist in the gene pool, transposable elements have to constantly infect novel hosts (Sawyer *et al*. 1987; Lawrence *et al*. 1992; Bichsel *et al*. 2010; Rankin *et al*. 2010; Wu *et al*. 2015; Park *et al*. 2021). Yet, there is at least one exception: a class of transposable elements called REPINs. REPINs are short (∼100 bp) nested palindromic sequences (**Figure 1**) that consist of two inverted REP (repetitive extragenic palindromic (Higgins *et al*. 1982)) sequences that can be present hundreds of times per genome (Bertels, Rainey 2011a). Most REPINs are symmetric where the 5’ REP sequences are identical to the 3’ REP sequences, with the occasional substitution (Bertels, Rainey 2011a; b). However, there are also asymmetric REPINs where the 5’ REP sequence differs from the 3’ REP sequence by a point deletion or insertion (Bertels, Rainey 2011a, 2023), which makes the analysis and detection of REPINs significantly more difficult (*e*.*g*., *Escherichia coli* REPINs). Isolated REP sequences, sometimes referred to as REP singlets, can also be found in the genome. These sequences are decaying remnants of REPINs that are no longer mobile (Bertels, Rainey 2011a). REPINs are non-autonomous mobile genetic elements, which means they require a RAYT (**R**EP **A**ssociated t**Y**rosine **T**ransposase) transposase gene (also referred to as tnpA_REP_) to replicate inside the genome (Nunvar *et al*. 2010; Bertels, Rainey 2011a; Ton-Hoang *et al*. 2012).

**Figure 1.**
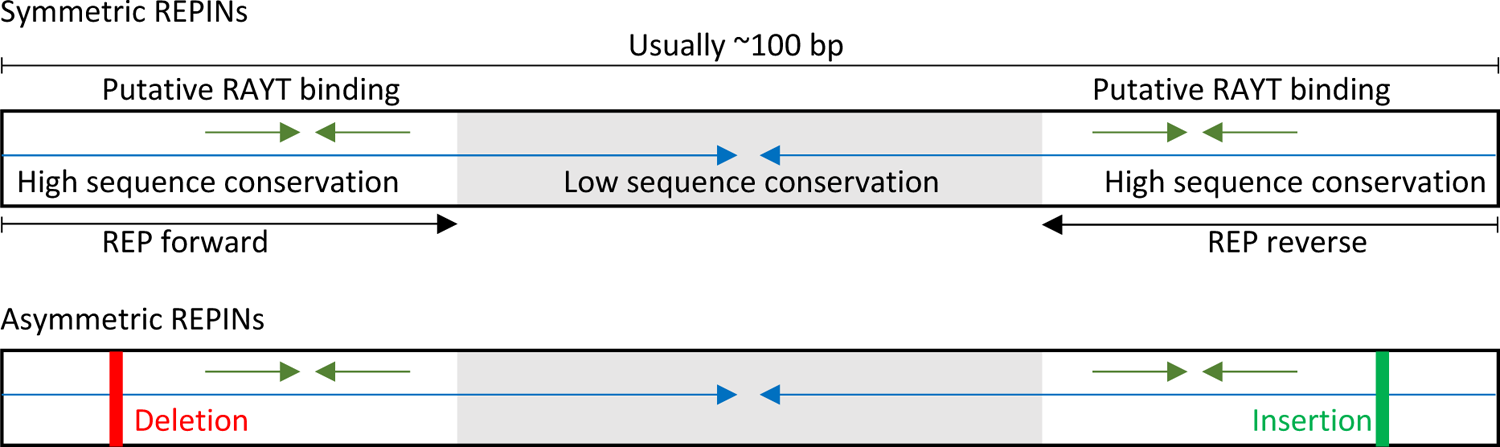
The structure of symmetric and asymmetric REPINs. A typical REPIN consists of two highly conserved regions at the 5’ and 3’ end (white), separated by a spacer region of lower sequence conservation (grey). The entire REPIN is an imperfect palindrome (blue arrows), which means it can form hairpin structures in single stranded DNA or RNA. Each 5’ and 3’ region contains a nested imperfect palindrome, which is referred to as REP (repetitive extragenic palindromic) sequence and has first been described in *Escherichia coli* (Higgins *et al*. 1982). REPINs can be either symmetric or asymmetric. Asymmetric REPINs have a deletion and a corresponding insertion in the highly conserved 5’ or 3’ end, which leads to “bubbles” in the hairpin structure. REPINs in for example *Pseudomonas fluorescens* SBW25 are symmetric while REPINs in *E. coli* are asymmetric. Asymmetric REPINs make analyses with RAREFAN more challenging. Figure adapted from (Bertels, Rainey 2023), which is licensed under <u>CC BY 4.0</u>.

Within a genome, each REPIN population is usually only associated with a single RAYT gene. Hence, RAYT genes occur only in single copies per genome and do not copy themselves, unlike for example insertion sequences where often multiple identical sequences are present inside the genome. Unlike insertion sequences RAYT genes are almost exclusively inherited vertically, meaning they are host-beneficial transposases that have been coopted by the host (Bertels, Gallie, *et al*. 2017; Bertels, Rainey 2023). The fact that REPINs and their corresponding RAYT genes are confined to a single bacterial lineage makes them unique, in comparison to all other parasitic mobile genetic elements in bacterial genomes (Bertels, Rainey 2023).

Of a total of five different RAYT families, there are only two RAYT families that are associated with repetitive sequences such as REPIN or REP sequences: Group 2 and Group 3 RAYTs (Bertels, Gallie, *et al*. 2017). Group 2 RAYTs are present in most Enterobacteria and usually occur only once per genome associated with a single REPIN population. In contrast, Group 3 RAYTs are found in most *Pseudomonas* species and are usually present in multiple divergent copies per genome, each copy associated with a specific REPIN population (Bertels, Gallie, *et al*. 2017).

REPINs and their corresponding RAYT genes occur exclusively in bacterial genomes and are absent in eukaryotic or archaeal genomes (Bertels, Gallie, *et al*. 2017; Bertels, Rainey 2023). Within bacterial genomes REPINs and RAYTs have been evolving in single bacterial lineages for millions of years (Bertels, Gallie, *et al*. 2017). The long term persistence of REPINs in single bacterial lineages can also be observed when analyzing REPIN populations (Bertels, Gokhale, *et al*. 2017; Bertels, Rainey 2023).

Parasitic insertion sequences usually occur in identical copies in bacterial genomes, reflecting the fact that insertion sequences persist only briefly before they are eradicated from the genome or kill their host (Park *et al*. 2021). REPINs in contrast are only conserved at the ends of the sequence (presumably due to selection for function), the rest of the sequence is highly variable and only the hairpin structure is conserved (Bertels, Rainey 2011a). The sequence variability of REPINs within the same genome reflects their long-term persistence in single bacterial lineages (Bertels, Rainey 2023). REPINs cannot simply reinfect another bacterial lineage since they rely for mobility on their corresponding RAYT, which itself is immobile.

RAYTs and REPINs are distinct from typical parasitic insertion sequences, yet we know very little about their evolution or biology. Currently, it is completely unclear what kind of beneficial function maintains REPINs and RAYTs as well as their association with each other. The reason for our lack of knowledge is not because REPINs and RAYTs are rare. They are ubiquitously found in many important and well-studied model bacteria such as Enterobacteria, Pseudomonads, Neisseriads and Xanthomonads. Microbial molecular biologists presumably encounter REPINs quite frequently. However, connecting the presence or absence of REPINs/RAYTs with phenotypes is difficult if we do not know when it is a REPIN that is present close to a gene of interest or a different type or repeat sequence. Even if the scientist knows about the presence of a REPIN, it is also important to know whether a corresponding RAYT is present (Bertels, Rainey 2023).

The identification of REPIN populations and their corresponding RAYTs can be rather cumbersome if done from scratch. This is particularly true if the microbial molecular biologist is not aware of all the details of REPIN and RAYT biology. Identifying REPINs starts with an analysis of short repetitive sequences in a genome. If there are excessively abundant short sequences present in the genome, the distribution of these sequences is then analyzed. If these sequences are exclusively identical tandem repeats without sequence variation, and present in only one or two loci in the genome, then these sequences are probably part of a CRISPR array and not REPINs. If the sequences are distributed across the genome, highly diverse and often present as inverted repeats then it is likely that the repeats are indeed REPINs.

Here, we present RAREFAN (RAYT/REPIN Finder and Analyzer), a webservice that automates the identification of REPINs and their corresponding RAYTs. RAREFAN is publicly accessible atu http://rarefan.evolbio.mpg.de and identifies REPIN populations and RAYTs inside a set of bacterial genomes. RAREFAN also generates graphs to visualize the population dynamics of REPINs, and assigns RAYT genes to their corresponding REPIN groups. Here we will demonstrate RAREFAN’s functionality by analyzing REPIN-RAYT systems in the bacterial species *Stenotrophomonas maltophilia*.

## Methods

### Identification of REPs and REPINs

The algorithm to determine REP sequence groups has been described in previous papers and is now slightly improved (Bertels, Rainey 2011a, 2023; Bertels, Gokhale, *et al*. 2017). The main difference between the implementations is that RAREFAN automatically links REPs/REPIN populations to RAYT genes, which was not possible previously.

The algorithm starts by extracting all N bp (21 bp by default) long seed sequences that occur more than M times (55 by default) from the reference genome (**Figure 2B**). All sequences occurring within the reference genome at least once within 15 bp of each other are then grouped together into n REP sequence groups (numbered 0-[n-1]). The most common sequence in each group, named REP seed sequence, is used for further analyses in each input genome.

**Figure 2.**
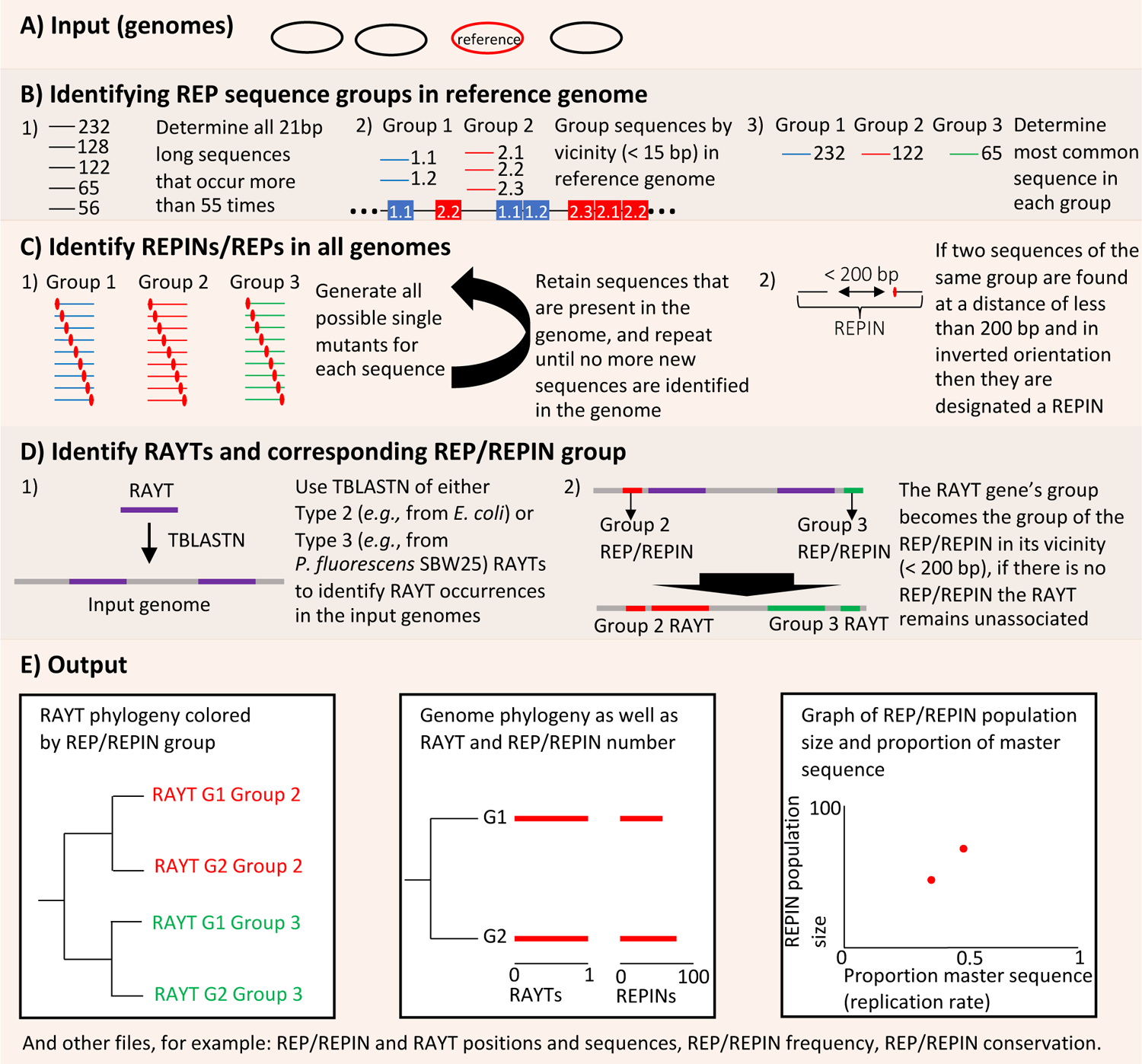
RAREFAN workflow. (**a**) By default RAREFAN requires the user to supply input sequences containing RAYTs and REPINs. These are fully sequenced and complete genomes. (**b**) RAREFAN then identifies seed sequence groups (potential REP sequences) in the reference genome by first isolating all 21 bp (adjustable parameter) long sequences that occur more than 55 times (adjustable parameter) in the reference genome. It is likely that a large number of these sequences belong to the same REPIN sequence type since the conserved part of REPINs is longer than 21 bp. Hence, we grouped all sequences together that occur within 15 bp (adjustable parameter) of each other anywhere in the genome. All further analyses are performed only with the most common sequence in each sequence group. This sequence will be called the seed sequence. (**c**) The occurrences of the seed and mutated seed sequences are identified in all submitted genomes. If a mutated seed sequence is identified in a genome, then all single mutants of that seed sequence are searched recursively in the same genome. All identified sequences that occur within 130 bp in inverted orientation of each other are designated REPINs. All other identified seed sequences and mutated seed sequences are REP singlets. (**d**) TBLASTN is used to identify RAYT homologs (e-Value < 1e-30, adjustable parameter) of either *E. coli* (Group 2 RAYT) or from *P. fluorescens* SBW25 (Group 3 RAYT) across all submitted genomes. If a RAYT homolog is in the vicinity (default < 200 bp, adjustable parameter) of a previously identified REPIN or REP singlet, then this RAYT is designated as associated with this REPIN group. (**e**) The first graph contains a RAYT phylogeny computed from a nucleotide alignment of all identified RAYT genes. The RAYT phylogeny indicates what RAYTs are associated with what REPIN populations (largest sequence cluster calculated with MCL) via colour coding. In a second graph the abundance of each REPIN population and RAYT copy number are displayed on a genome phylogeny. In the last graph REPIN population sizes are plotted in relation to the proportion of master sequences. Master sequences are the most abundant REPIN in each population. RAREFAN also generates various files containing, for example, REP, REPIN, or RAYT sequences and their positions in the query genomes.

In the next step all possible point mutants of the seed sequences are generated and searched for in the genome (**Figure 2C**). If a sequence is found in the genome, then all possible point mutants are generated for this sequence as well and searched against the genome and so on until no more sequences can be identified. Once all sequences related to the seed sequence are found, RAREFAN determines whether the sequences form REPINs. Two sequences form a REPIN when they are located within 130 bp of each other in inverted orientation.

REP and REPIN sequences of the same type form REPIN populations. A REPIN population is defined as the largest coherent sequence cluster. To identify the largest sequence cluster MCL is applied to a network of REP/REPIN sequences where all sequences that differ by exactly one nucleotide are connected using an inflation parameter of 1.2 (Van Dongen 2000). The clustering results are stored in a file ending in .mcl. The sequences of the largest REPIN population (excluding REP singlets) are stored in a file ending in largestCluster.nodes. The largest REPIN populations are shown in the REPIN population plot and the master sequence correlation plot (**Figure 4**).

### Identification of RAYTs

RAYTs are identified using TBLASTN (Camacho *et al*. 2009) with either a protein sequence provided by the user, a Group 2 RAYT from *E. coli* (yafM, Uniprot accession Q47152) or a Group 3 RAYT from *P. fluorescens* SBW25 (yafM, Uniprot accession C3JZZ6). The presence of RAYTs in the vicinity (default maximum of 200 bp) of a particular REPIN can be used to establish the association between the RAYT gene and a REPIN group (**Figure 2D**). If a REP sequence or a REPIN of a particular group occur within 200 bp (by default) of a RAYT gene then the RAYT gene is linked to the REP/REPIN group. REP/REPIN associations are stored in the file repin_rayt_association.txt. REPIN or REP sequences are almost always present in the extragenic spaces of the RAYT and this linkage is consistent across the RAYT phylogeny (as shown in **Figure 5**). However, what causes the REP/REPIN group to be linked with the RAYT gene is unclear.

### Implementation

RAREFAN is a modular webservice. It consists of a web frontend written in the Python programming language (Van Rossum, Drake Jr 1995) using the Flask framework (Grinberg 2018), a Java (Arnold *et al*. 2005) backend for genomic sequence analysis and an R (R Core Team 2016) Shiny app (RStudio, Inc 2013) for data visualization. The software is developed and tested on the Debian GNU/Linux operating system (Kleinmann *et al*. 2021). All components are released under the MIT opensource license (Initiative 2021) and can be obtained from our public GitHub repository at https://github.com/mpievolbio-scicomp/rarefan.

The public RAREFAN instance at http://rarefan.evolbio.mpg.de runs on a virtual cloud server with four single-threaded CPUs and 16GB of shared memory provided and maintained by the Gesellschaft für Wissenschaftliche Datenverarbeitung Göttingen (GWDG) and running the Debian GNU/Linux Operating System (Kleinmann *et al*. 2021).

The Java backend drives the sequence analysis. It makes system calls to TBLASTN (Altschul *et al*. 1990) to identify RAYT homologs and to MCL (Van Dongen 2000) for clustering REPIN sequences in order to determine REPIN populations.

Jobs submitted through the web server are queued and executed as soon as the required resources become available. Users are informed about the status of their jobs. After job completion, the user can trigger the R Shiny app to visualize the results.

The Java backend can also be run locally *via* the command line interface (available for download at https://github.com/mpievolbio-scicomp/rarefan/releases).

### Usage of the webservice

The front page of our webservice allows users to upload their bacterial genomes in FASTA (.fas) format (**Figure 2A**). Optionally, users may also provide RAYT protein FASTA sequences (.faa) or phylogenies in NEWICK (.nwk) format. After successful completion of the upload process, the user fills out a web form to specify the parameters of the algorithm:

- Reference sequence: Which of the uploaded genome sequences will be designated as reference genome (see below for explanations). Defaults to the first uploaded filename in alphabetical order.
- Query RAYT: The RAYT gene that is used to identify homologous RAYTs in the query genomes. If the user does not provide a protein sequence file then the user can choose one of two RAYT sequences (one from Group 2 and one from Group 3 RAYTs (Bertels, Gallie, *et al*. 2017)) as RAYT query.
- Tree file: A phylogenetic tree of the reference genomes that can be provided by the user, otherwise the tree will be calculated using andi (Haubold *et al*. 2015).
- Minimum seed sequence frequency: Lower limit on seed sequence frequency in the reference genome to be considered as a REP candidate. Default is 55.
- Seed sequence length: The seed sequence length (in base pairs) is used to identify REPIN candidates from the input genomes. Default is 21 bp.
- Distance group seeds: The maximum distance between a single occurrence of short repetitive sequences to still be sorted into the same sequence group.
- REPIN-RAYT association distance: The maximum distance at which a REP sequence can be located from a RAYT gene to be linked to that RAYT gene. Default is 200 bp.
- e-value cut-off: Alignment e-value cut-off for identifying RAYT homologs with TBLASTN. Default is 1e-30.
- Analyse REPINs: Ticked REPINs will be analysed (two inverted REP sequences found at a distance of less than 130 bp), if not ticked only short repetitive 21 bp long sequence will be analysed.
- User email (optional): If provided, then the user will be notified by email upon run completion.

The job is then ready for submission to the job queue. Upon job completion, links to browse and to download the results, as well as a link to a visualization dashboard are provided. If a job runs for a long time then users may also come back to RAREFAN at a later time, query their job status and eventually retrieve their results by entering the run ID into the search field at http://rarefan.evolbio.mpg.de/results. Relevant links and the run ID are communicated either on the status site or by email if the user provided their email address during run configuration. Runs are automatically deleted from the server after six months.

### Visualizations

For each REPIN-RAYT group summary plots are generated. These include plots showing the RAYT phylogeny (calculated from a nucleotide alignment using MUSCLE (Edgar 2004) and PHYML (Guindon *et al*. 2010) to generate a phylogeny), REPIN population sizes in relation to the genome phylogeny (provided by the user or if not provided calculated by andi (Haubold *et al*. 2015)) as well as the proportion of master sequences (most common REPIN in a REPIN population) in relation to REPIN population size (**Figure 2E**).

### Other outputs

Identified REPINs, REP singlets as well as RAYTs are written to FASTA formatted sequence files and to tab formatted annotation files that can be read with the Artemis genome browser (Rutherford *et al*. 2000). The REPIN-RAYT associations as well as the number of RAYT copies per genome are written to tabular data files. A detailed description of all output files is provided in the manual (http://rarefan.evolbio.mpg.de/manual) and in the file “readme.md” in the output directory.

### Sequence analysis and annotation

For verification of RAREFAN results, REPIN-RAYT-systems were analysed in their corresponding genomes using Geneious Prime version 2022.2.2 (Kearse *et al*. 2012). Nucleotide sequences and positions of REP singlets, REPINs, and RAYTs were extracted from output files generated by RAREFAN and mapped in the relevant *S. maltophilia* genome. Complete RAREFAN data used for analysis can be accessed by using the run IDs listed in **Table 1**.

**Table 1.**
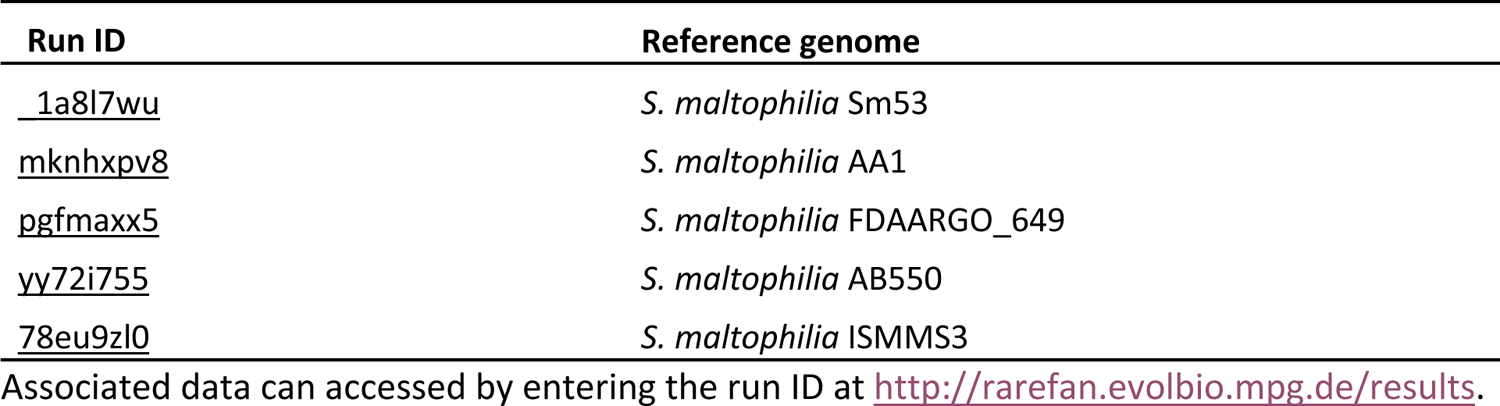
RAREFAN IDs linking to the raw data of the presented analyses.

## Results

RAREFAN can identify REPINs and their corresponding RAYTs in a set of fully sequenced bacterial genomes. The RAREFAN algorithm has been used in previous analyses to identify and characterize REPINs and RAYTs in Pseudomonads (Bertels, Rainey 2011a, 2023), Neisseriads (Bertels, Rainey 2023), and Enterobacteria (Bertels, Gallie, *et al*. 2017; Park *et al*. 2021). To demonstrate RAREFAN’s capabilities, we are presenting an analysis of 49 strains belonging to the opportunistic pathogen *S. maltophilia*.

*S. maltophilia* strains contain Group 3 RAYTs, which are also commonly found in plant-associated *Pseudomonas* species such as *P. fluorescens* or *P. syringae* (Bertels, Rainey 2011a, 2023). Similar to Group 3 RAYTs in other species, *S. maltophilia* contains multiple REPIN-RAYT systems per genome. Group 2 RAYTs, in contrast, tend to contain only one REPIN-RAYT system per genome (Bertels, Rainey 2023).

### *Nine different REPIN-RAYT systems in* S. maltophilia

REPIN-RAYT systems in *S. maltophilia* are surprisingly diverse compared to other species. For example, *Pseudomonas chlororaphis* contains three separate REPIN populations that are present in all *P. chlororaphis* strains, each associated with its cognate RAYT gene (Bertels, Rainey 2023). *S. maltophilia*, in contrast, contains only one REPIN-RAYT system that is present across almost the entire species (green clade in **Figure 3**), and at least eight REPIN-RAYT systems that are present in subsets of strains (nine clades in **Figure 5**).

**Figure 3.**
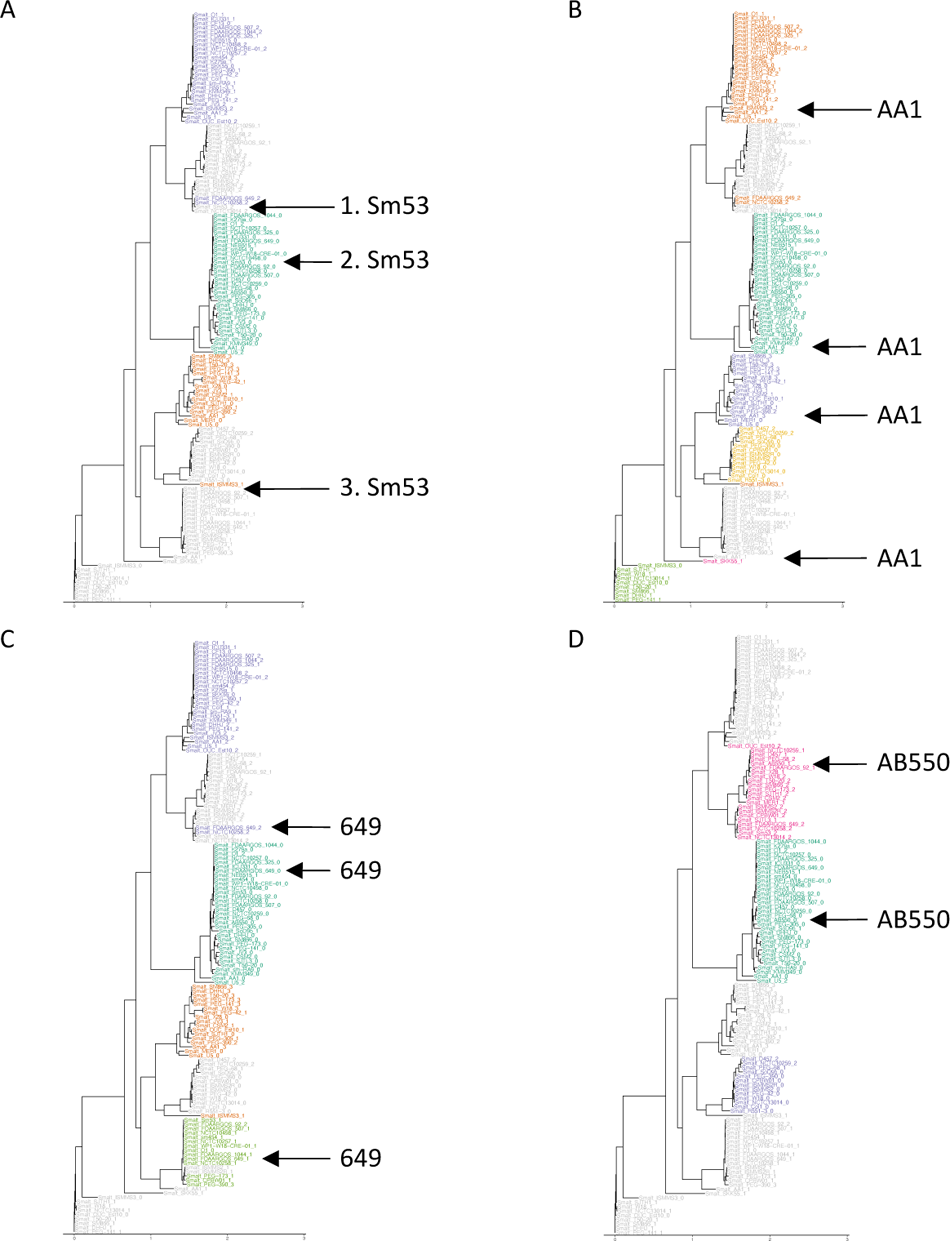
Phylogenetic trees built from RAYT genes extracted from *S. maltophilia* genomes. RAYT genes are coloured by RAREFAN according to their association with REPIN populations in the reference genome. If a REPIN population of a query genome is not present in the reference genome, then the REPIN population cannot be identified in the query genome and the corresponding RAYT gene cannot be linked and is coloured in grey. The four panels **A-D** show phylogenies for four different reference strains. *S. maltophilia* strains Sm53, AA1, 649 and AB550 were used in panels **A** to **D**, respectively. Locations of a reference strain’s RAYT genes in the tree are indicated by arrows. An association between almost all RAYTs and REPIN populations could be made by using four different reference genomes. Most of the RAYT genes are coloured (associated with a REPIN group) in at least one of the trees. The three numbered RAYT genes from the Sm53 RAREFAN run are referenced in the text.

The patchy presence-absence pattern of REPIN-RAYT systems in *S. maltophilia*, makes the dataset quite challenging to analyse. If a REPIN population is not present in the reference strain then RAREFAN will not be able to detect it in any other strain. Yet, it is possible to detect RAYT genes in all strains of a species independent of the reference strain selection. RAYT genes that are not associated with a REPIN population are displayed in grey (**Figure 3A**). While these RAYT genes are not associated with REPIN populations detected in the reference strain, they might still be associated with a yet unidentified REPIN type present in the genome the unassociated RAYT gene is located in.

In order to identify all REPIN populations across a species, multiple RAREFAN runs with different reference strains should be performed. The RAREFAN web interface supports re-launching a given job with modified parameters. To identify as many different REPIN-RAYT systems as possible in each subsequent run the reference should be set to a genome that contains RAYTs that were not associated with a REPIN population previously (*i*.*e*., genomes containing grey RAYTs in **Figure 3**). However, this strategy may also fail when the REPIN population size falls below the RAREFAN seed sequence frequency threshold. In that case reducing the frequency threshold will be more productive.

For example, *S. maltophilia* Sm53 contains three RAYTs only one of which is associated with a REPIN population (RAYT genes indicated in **Figure 3A**). However, the remaining two RAYTs are indeed associated with a REPIN population, but these REPIN populations are too small to be detected in *S. maltophilia* Sm53 (the seed sequence frequency threshold is set to 55 by default). In other *S. maltophilia* strains the REPIN populations are large enough to exceed the threshold. For example, if *S. maltophilia* AB550 is set as reference, RAYT number 1 from Sm53 (**Figure 3A**) is associated with the pink REPIN population (**Figure 3D**). If *S. maltophilia* 649 is set as reference RAYT number 3 from Sm53 (**Figure 3A**) is associated with the light green REPIN population (**Figure 3C**). RAYTs from the bottom clade are only associated with REPIN populations when *S. maltophilia* AA1 is chosen as reference (**Figure 3B**). While lower thresholds can guarantee that all REPINs will be identified in the genome, the number of sequence groups that are not REPINs quickly explodes. This is especially true for genomes that contain large numbers of mobile genetic elements or CRISPRs (Bertels, Rainey 2023).

### RAREFAN visualizes REPIN population size and potential replication rate

The RAREFAN webserver visualizes REPIN population size and RAYT numbers in barplots. Barplots are ordered by the phylogenetic relationship of the submitted bacterial strains using ggtree (Yu *et al*. 2018). RAREFAN detects three populations when *S. maltophilia* Sm53 is selected as reference strain (**Figure 3A**). The largest REPIN population (calculated by MCL from all REPINs of that type) has a corresponding RAYT gene in almost all strains (first barplot in **Figure 4A**) and most REPIN populations contain more than 100 REPINs (second barplot in **Figure 4A**). The second largest REPIN population in Sm53 (orange population in **Figure 4B**) is significantly smaller and contains no more than 61 REPINs in any strain and most strains do not contain a corresponding RAYT for this population.

**Figure 4.**
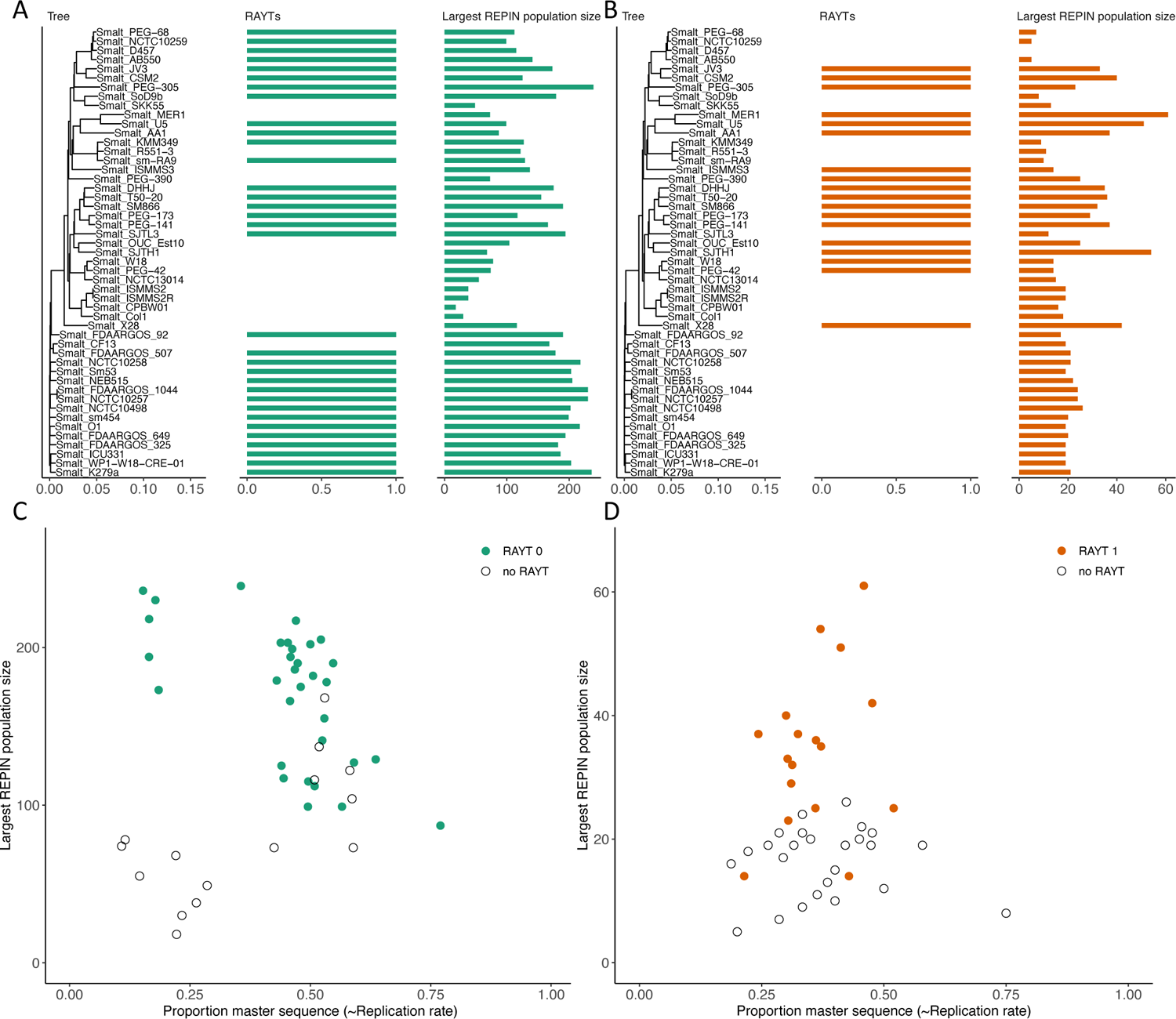
REPIN population sizes and conservation. The plots show two REPIN populations and their associated RAYTs that were identified in *S. maltophilia* using *S. maltophilia* Sm53 as reference. (**A**) The phylogenetic tree on the left side is a whole genome phylogeny generated by andi (Haubold *et al*. 2015). Shown on the right are REPIN population sizes (which is the largest REPIN cluster calculated by MCL) and the number of associated RAYTs sorted by the genome phylogeny. The green REPIN populations and associated RAYTs are present in most strains in high abundance (maximum of 239 occurrences in *S. maltophilia* K279a, left panel). (**B**) The orange population in contrast is present in much lower numbers (maximum of 61 occurrences in *S. maltophilia* MER1, right panel). Note, REPIN populations are assigned consistent colours based on their abundance in the reference genome. For example, the most abundant REPIN population in the reference is always coloured in green, and the second most abundant population is always coloured in orange. (**C and D**) Proportion of master sequence in *S. maltophilia* REPIN populations. The master sequence in a REPIN population is the most common REPIN sequence. At equilibrium the higher the proportion of the master sequence in the population the higher the replication rate (Bertels, Gokhale, *et al*. 2017). The presence and absence of an associated RAYT is also indicated by the colours of the dots. Empty circles indicate that the REPIN population is not associated with a RAYT gene in that genome.

RAREFAN also provides information on REPIN replication rate (**Figure 4C** and **D**). REPIN replication rate can be estimated by dividing the number of the most common REPIN sequence (master sequence) by the REPIN population size if the population is in mutation selection balance (Bertels, Gokhale, *et al*. 2017). If a REPIN replicates very fast most of the population will consist of identical sequences because mutations do not have enough time to accumulate between replication events. Hence, the proportion of master sequences will be high in populations that have a high replication rate. Transposable elements such as insertion sequences consist almost exclusively of identical master sequences because the time between replication events is not sufficient to accumulate mutations and because quick extinction of the element usually prevents the accumulation of mutations after replication (Park *et al*. 2021; Bertels, Rainey 2023). Sequence diversity of REPIN populations in contrast is much higher suggesting that REPINs replicate slowly and persist for long periods of time.

In *S. maltophilia* the proportion of master sequences in the population does not seem to correlate well with REPIN population size, both in the green and the orange population (**Figures 4C and D**). Similar observations have been made in *P. chlororaphis* (Bertels, Rainey 2023), and may suggest that an increase in population size is not caused by an increase in replication rate. Population size is likely to be more strongly affected by other factors such as the loss of the corresponding RAYT gene, which leads to the decay of the REPIN population. One could even speculate that high REPIN replication rates are more likely to lead to the eventual demise of the population due to the negative fitness effect of high replication rates on the host (Park *et al*. 2021; Bertels, Rainey 2023).

The presence of RAYTs and the size of the corresponding REPIN population do correlate surprisingly well (**Figure 4A** and **B**, p-value = 0.008 of a linear model of independent contrasts (Felsenstein 1985) of green RAYT and REPIN number, p-value = 0.003 for orange REPIN populations). Green RAYTs are absent from an entire *S. maltophilia* clade (middle of **Figure 4A**). This clade has also lost a significant amount of green REPINs, and the proportion of the master sequences in these populations is low (**Figure 4C**). Similarly, genomes without orange RAYTs have smaller REPIN populations in the orange population than genomes with the corresponding RAYT (**Figure 4D**). A similar observation has been made previously in *E. coli, P. chlororaphis, N. meningitidis* and *N. gonorrhoeae* where the loss of the RAYT gene is followed by a decay of the associated REPIN population (Park *et al*. 2021; Bertels, Rainey 2023).

### Linking REPIN populations with RAYT genes can be challenging

Unfortunately, RAREFAN is not always able to link the correct REPIN population with the correct RAYT gene. In some RAREFAN runs, associations between RAYTs and REPINs are not monophyletic, as for example the red RAYT clade in **Figure 3A**. However, the same clade of RAYTs is uniformly coloured in yellow in **Figure 3D**, suggesting that the entire RAYT clade is associated with the same REPIN group.

An analysis of all REPIN groups that were identified by RAREFAN across four different RAREFAN runs (**Table 1**, one additional analysis was performed with ISMMS3) showed that there are a total of nine different REPIN groups, each defined by an individual central palindrome (**Table 2**). Each REPIN group is associated with a monophyletic RAYT group (**Figure 5**). Only a single RAYT is not associated with a REPIN population (ISMMS3_1).

**Figure 5.**
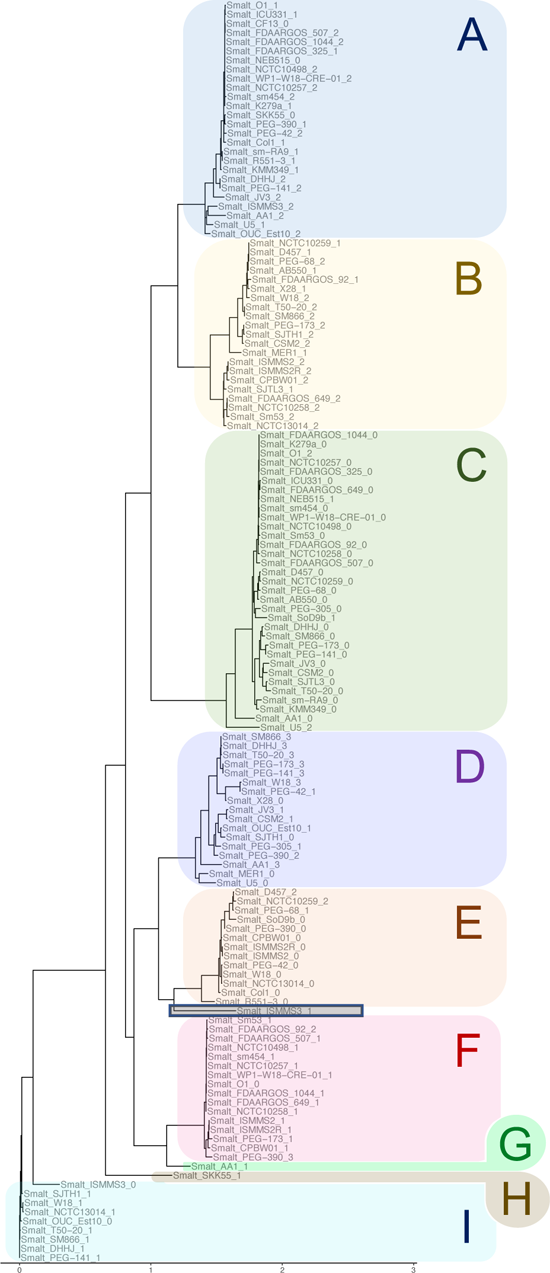
Phylogeny of RAYT genes and their associated REPINs. The tree shows RAYT genes from 49 *S. maltophilia* strains. Colours of clades A-I are assigned according to their association with a REPIN found within 200 bp of the RAYT gene (see **Table 2**). Except for a single RAYT gene ISMMS3_1 (grey box), which could not be linked to a REPIN population.

**Table 2.**
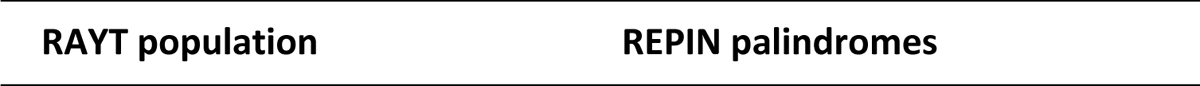
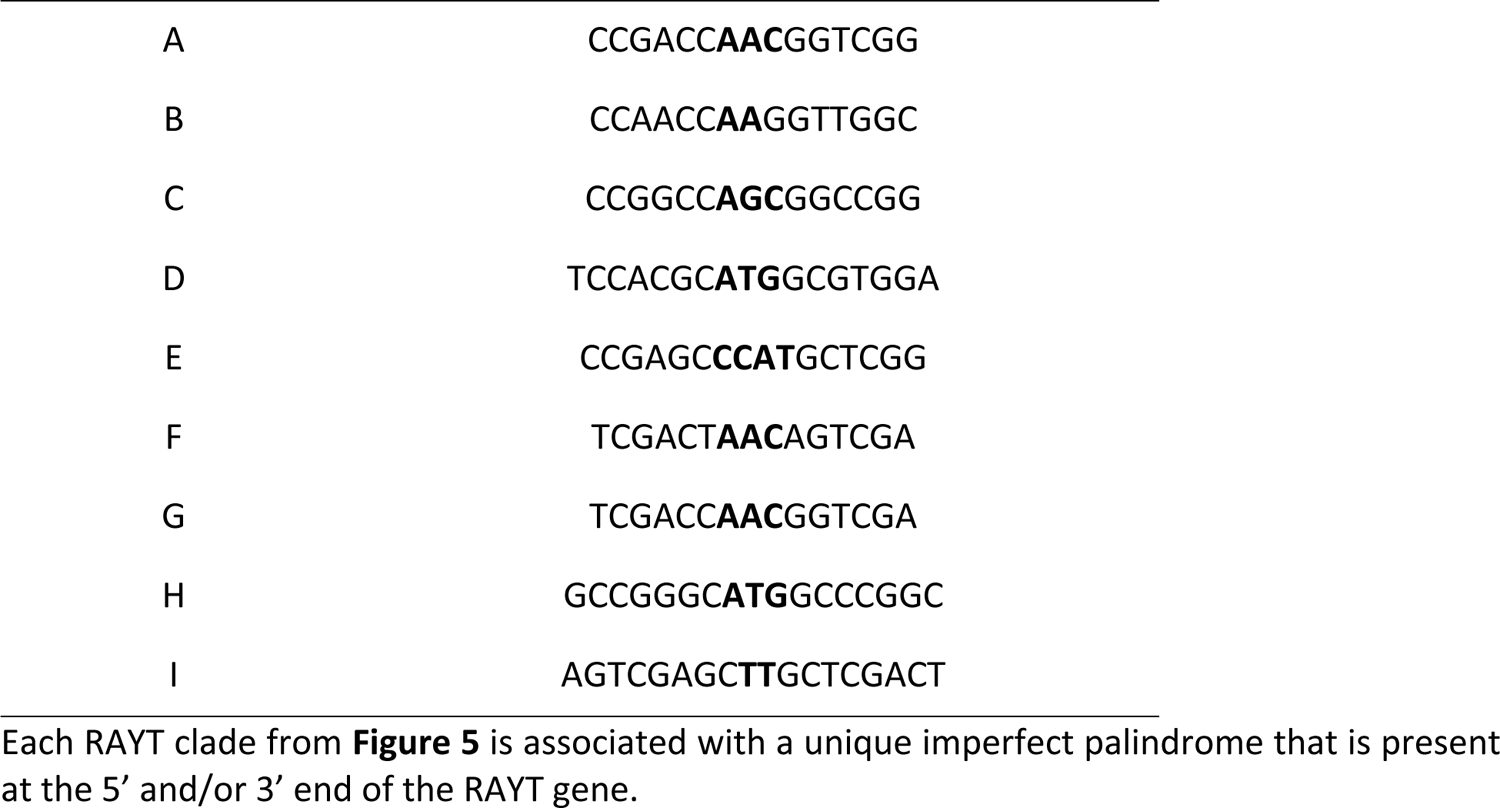
REPIN palindromes associated with each RAYT clade.

RAREFAN could not link a REPIN to the RAYT gene ISMMS3_1 (**Figure 5**, grey box). While there is a sequence that resembles the A palindrome as well as variants of the C palindrome flanking both sides of the RAYT gene (**Supplementary Figure 2**), none of the sequences formed REPIN populations large enough to be identified by RAREFAN. Presumably the RAYT ISMMS3_1, which is only present in a single *S. maltophilia* strain, is at the early stages of establishing a REPIN population, and the REPIN population has not spread to a considerable size yet.

If the maximum REPIN-RAYT distance parameter is too small then RAREFAN will also fail to correctly link REPINs and RAYTs. For example, when the maximum REPIN-RAYT distance parameter is set to 130bp there are two cases where RAREFAN fails to link RAYT genes with REPINs (ISSMS2_ and ISSMS2R_1, **Supplementary Figure 1 D** and **E**). When the parameter is set to a distance of 200 bp (default RAREFAN setting), RAREFAN correctly links these REPINs to the RAYT gene.

In three cases the wrong REPIN population was linked to a RAYT gene. In our dataset this can happen when RAYTs are flanked by seed sequences from two different REPIN populations **(Supplementary Figure 1 A-C)**. A single REP sequence from the “wrong” (non-monophyletic RAYT) clade occurs together with multiple REP or REPIN sequences from the “right” (monophyletic in a different RAREFAN run) clade. REPINs are linked to the “wrong” RAYT when the correct REPIN population is absent in the chosen reference genome. This problem can be alleviated by performing analyses with multiple reference genomes and comparing the results.

### REPIN groups may be lost when the seed distance is too large

The seed distance parameter determines whether two highly abundant sequences are sorted into the same or different REPIN groups (**Figure 2B**). If two REPINs from two different groups occur next to each other, at a distance of less than the seed distance parameter, then the two seeds are erroneously sorted into the same group. If two different REPIN groups are sorted into the same group then one of the groups will be ignored by RAREFAN, because only the most abundant seed in each group will be used to identify REPINs.

A manual analysis (*e*.*g*., multiple sequence alignment) of sequences in the groupSeedSequences folder of the RAREFAN output can identify erroneously merged REPIN groups. In *S. maltophilia*, groups are separated well when the distance parameter is set to 15 bp and Sm53 is used as a reference. When the parameter is set to 30 bp instead, one of the REPIN groups will be missed by RAREFAN.

A small seed distance parameter will separate seed sequences belonging to the same REPIN group into different groups. Hence, RAREFAN will analyse the same REPIN group multiple times. While this will lead to increased RAREFAN runtimes, these errors, are easy to spot, because (1) the same RAYT gene will be associated to multiple REPIN groups, (2) the central palindrome between the group is identical and (3) the master sequence between the groups will be very similar.

### Closely related REPIN groups may be merged into a single group by RAREFAN

Incorrect merging of REPIN groups can occur when two REPIN groups are closely related. We identified merged REPIN groups in *S. maltophilia* because RAREFAN linked some REPIN groups with two RAYT genes in the same genome (**Figure 6A**). While REPIN groups linked to two RAYTs have been observed before in *Neisseria meningitidis* (Bertels, Rainey 2023), it is particularly unusual in *S. maltophilia* due to some key differences between REPIN-RAYT in the two bacterial species. First, *N. meningitidis* contains Group 2 RAYTs and *S. maltophilia* only contains Group 3 RAYTs (Bertels, Gallie, *et al*. 2017), two very divergent RAYT gene families. Second, RAYTs that are associated with the same REPIN group in *N. meningitidis* are almost identical, since they are copied by an insertion sequence *in trans* (Bertels, Rainey 2023), something that is not the case for *S. maltophilia*, where the two RAYTs are very distinct and quite distantly related from each other (green and red clade in **Figure 3A**, or clade A and C in **Figure 5**).

**Figure 6.**
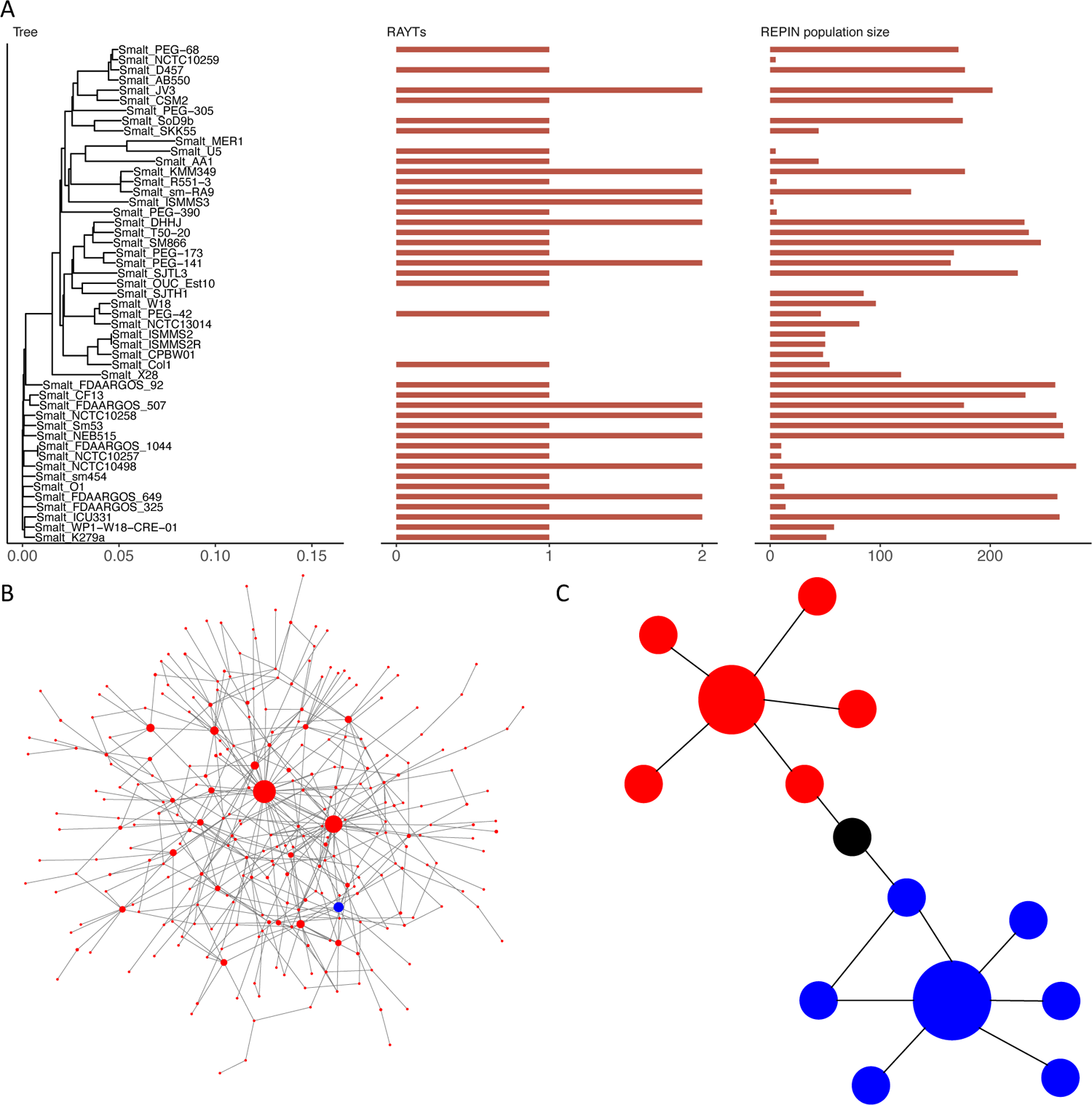
Closely related REPIN populations may be merged by RAREFAN. **(A)** REPIN group 2 identified in a *S. maltophilia* Sm53 RAREFAN run. The RAREFAN result suggests that REPIN group 2 is sometimes associated with two RAYTs. (**B**) A closer inspection of the data shows that group 2 is a combination of two different REPIN groups, the “real” group 2 and group 0. The network shown, visualizes all REP sequences identified as group 2. Nodes in the network represent 21 bp long REP sequences. Two nodes are connected if the sequences they represent differ by exactly one nucleotide. The node size indicates the abundance of the sequence in the genome. The blue node represents the most common group 2 sequence, occurring 65 times in the genome. The largest red node occurs 407 times in the genome and resembles a group 0 REP sequence. (**C**) Illustration of how small changes to a single sequence can connect two sequence clusters. The most common 21 bp long sequence in group 0 differs in only four positions from the most common 21 bp long sequence in group 2. There is a set of sequences that connects these two groups that only differ in exactly one position each (nodes connected by an edge), which passes through the black node. If there is such an unbroken path between REP sequences, then REPIN groups will be merged.

A closer inspection of all sequences identified in REPIN group 2 shows that it also contains sequences belonging to REPIN group 0 (palindromes linked to clade A and C in **Table 2**). The relationship between the sequences shows that there is a chain of sequences that all differ by at most a single nucleotide between the most abundant sequence in group 2 to the most abundant sequence in group 0 (**Figure 6B and C**). Hence, the reason group 0 and group 2 are merged is that they are too closely related to each other and hybrids of the two REPIN groups exist. Because sequence groups are built by identifying all related sequences in the genome recursively, closely related groups (the REPIN group 0 seed only differs in four nucleotides from the REPIN group 2 seed sequence) can be merged into a single REPIN group. REPIN population size and RAYT number are the sum of REPIN group 0 and 2. There are various possibilities to resolve this issue: (1) subtract sequences from group 0 (which does not contain group 2) from REPIN group 2; (2) use a different sequence seed from the group 2 seed collection in the seed sequence file (groupSeedSequences/Group_Smalt_Sm53_2.out); (3) sometimes it may be possible to rerun RAREFAN with a different reference strain where the issue does not occur; or (4) increase the length of the seed sequence.

### Performance

We measured the elapsed time for a complete RAREFAN run for three different species and for 5, 10, 20, and 40 genomes with randomly selected reference strains and the two query RAYTs (yafM_Ecoli and yafM_SBW25). For a given number N of submitted genomes of average sequence length L (in megabases), a RAREFAN run completes in approximately T = (8-10 seconds) * N * L on our moderate server hardware (4 CPU cores, 16 GB shared RAM) (**Supplementary Figure 3** and **4)**.

## Discussion

RAREFAN allows users to quickly detect REPIN populations and RAYT transposases inside bacterial genomes. It also links the RAYT transposase genes to the REPIN population it duplicates. These data help the user to study REPIN-RAYT dynamics in their strains of interest without a dedicated bioinformatician, and hence will render REPIN-RAYT systems widely accessible.

One limitation of RAREFAN is that REPINs can only be identified in genomes when they are symmetric (**Figure 1**). Symmetric REPINs have seed sequences that can morph into each other by a series of single substitutions (intermediate sequences need to be present in the genome). A REPIN consists of a 5’ and a 3’ REP sequence. If one of these REP sequences contains an insertion or deletion, which the other REP sequence does not contain then RAREFAN will not recognize the second repeat of the seed sequence. In this case, RAREFAN will not be able to identify REPINs but can still be used to analyze REP singlet populations. To date, the only asymmetric REPIN populations known to us are found in *E. coli*. However, it is likely that asymmetric REPINs also exist in other microbial species.

RAREFAN sometimes cannot correctly divide REPINs into REPIN groups. Either because REPINs from different groups occur in close proximity in the genome, an issue that can easily be solved by adjusting a RAREFAN parameter, or because two REPIN groups are very closely related (**Figure 6**). Unfortunately, RAREFAN is not able to automatically detect and resolve the assignment of closely related REPINs into groups yet. Hence it is advisable to manually check associations between REPIN groups and RAYT genes by analyzing the composition of REPIN groups.

In the future we aim to make RAREFAN even more versatile and easier to use by, for example, automatically integrating data from public databases such as GenBank, and creating a RAREFAN Galaxy workflow (Afgan *et al*. 2018).

RAREFAN makes the study of REPIN-RAYT systems more accessible to any biologist or bioinformatician interested in studying intragenomic sequence populations. Our tool will help understand the purpose and evolution of REPIN-RAYT systems in bacterial genomes.

## Acknowledgements

We would like to thank Prajwal Bharadwaj for assisting us with the sequence analysis and Jenna Gallie for valuable feedback on the manuscript.

## Supplementary Figures

**Supplementary Figure 1.**
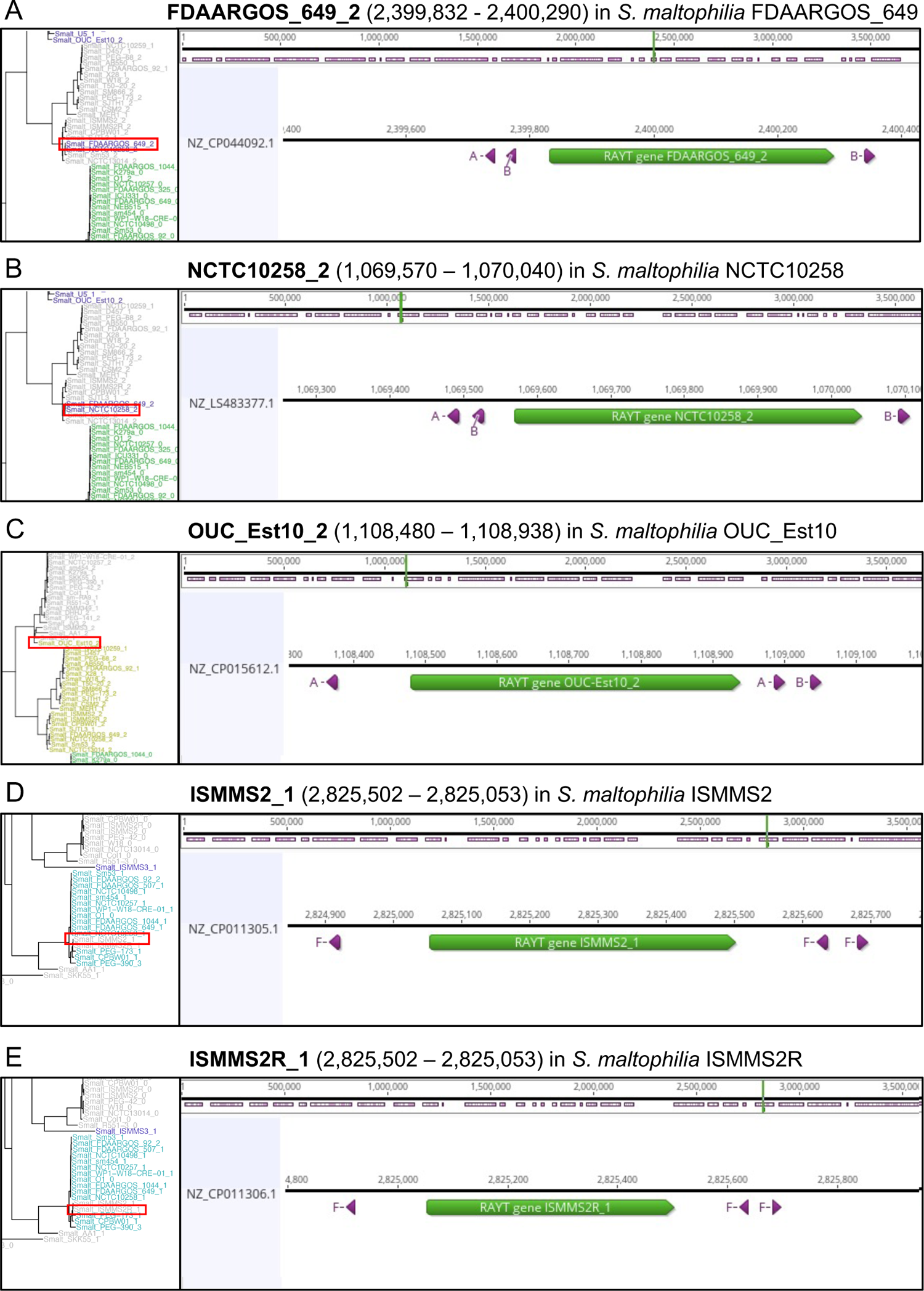
Sequence analysis shows REPIN groups are indeed associated with monophyletic RAYTs. Non-monophyletic or missing associations to REPIN populations identified by RAREFAN were investigated in the corresponding genomes using Geneious (Kearse *et al*. 2012). Red boxes mark the position of the atypical RAYT that is being analyzed in detail. Mapping of REPIN palindromes A-I (with zero mismatches) shows FDAARGOS_649_2 (A), NCTC10258_2 (B), and OUC_Est_2 (C) are linked to the wrong REPIN group because REP singlets that are ordinarily linked to a RAYT sister clade are found in close proximity to the RAYT. These wrong associations between REPIN and RAYT usually occur when the correct REPIN population is absent from the reference genome. ISMMS2R_1 (D) and ISMMS2_1 (E) are not linked to REPIN populations by RAREFAN when the maximum REPIN-RAYT distance parameter is set to 130 bp. The RAYTs are linked to the correct REPIN populations when the REPIN-RAYT distance parameter is set to 200 bp (default). Nucleotide sequences and positions were extracted from output files generated by RAREFAN. Complete genome sequences are available in NCBI Nucleotide Database using Accessions: (A) NZ_CP044092.1, (B) NZ_LS483377.1, (C) NZ_CP015612.1, (D) NZ_CP011306.1, (E) NZ_CP011305.1.

**Supplementary Figure 2.**
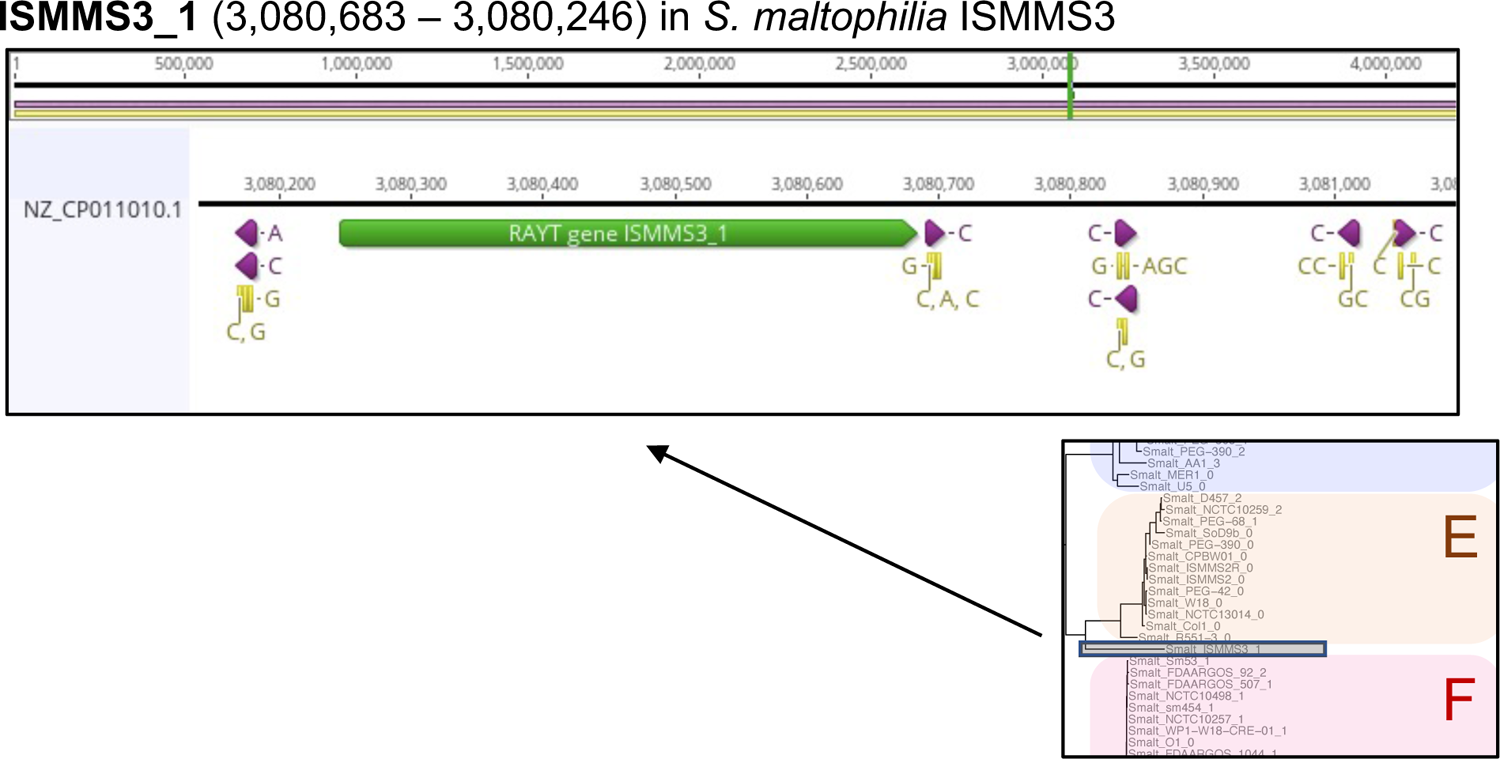
RAYT gene ISMMS3_1 cannot be linked to a REPIN population. The sequence of the RAYT gene ISMMS3_1 and its flanking sequences were analysed in Geneious (Kearse *et al*. 2012). The inset shows the location of ISMMS3_1 in the RAYT phylogeny (grey box). When mapping all of the identified palindromes to the RAYT region and allowing up to four mismatches (yellow annotations), various mutants of palindrome C were found in close proximity of the RAYT gene. However, we could not identify a corresponding REPIN population, which may indicate that the population has not yet expanded in the genome.

**Supplementary Figure 3.**
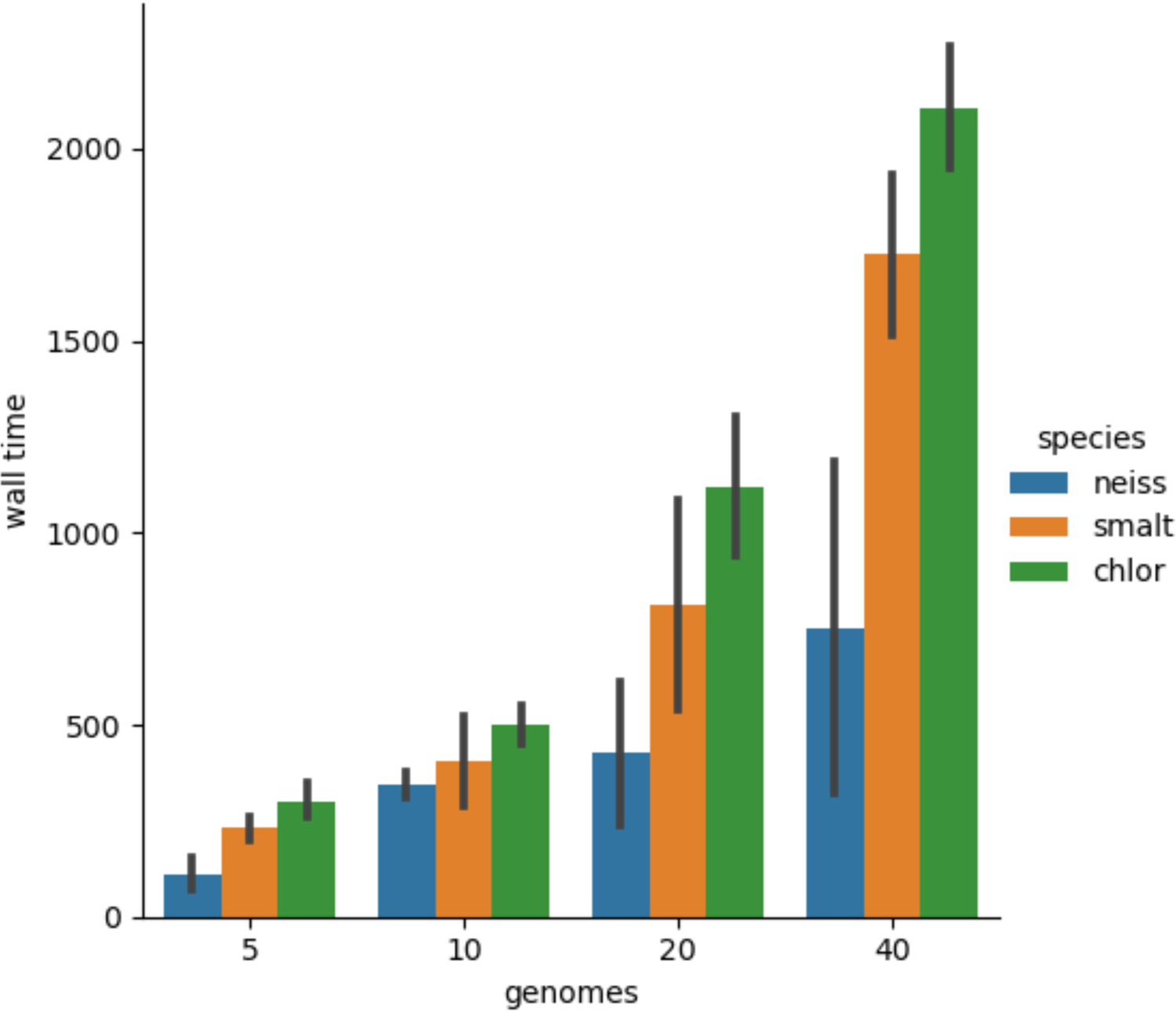
Average time (in seconds) it takes RAREFAN to complete for different genome numbers from three bacterial species (*N. meningitidis, N. gonorrhoeae, S. maltophilia, Pseudomonas chlororaphis*). Black bars indicate the 95% CI across four runs, where two runs share the same query RAYT. For each run reference and query strains were randomly selected. All measurements were performed on 4CPU cores with 16 GB of shared memory.

**Supplementary Figure 4.**
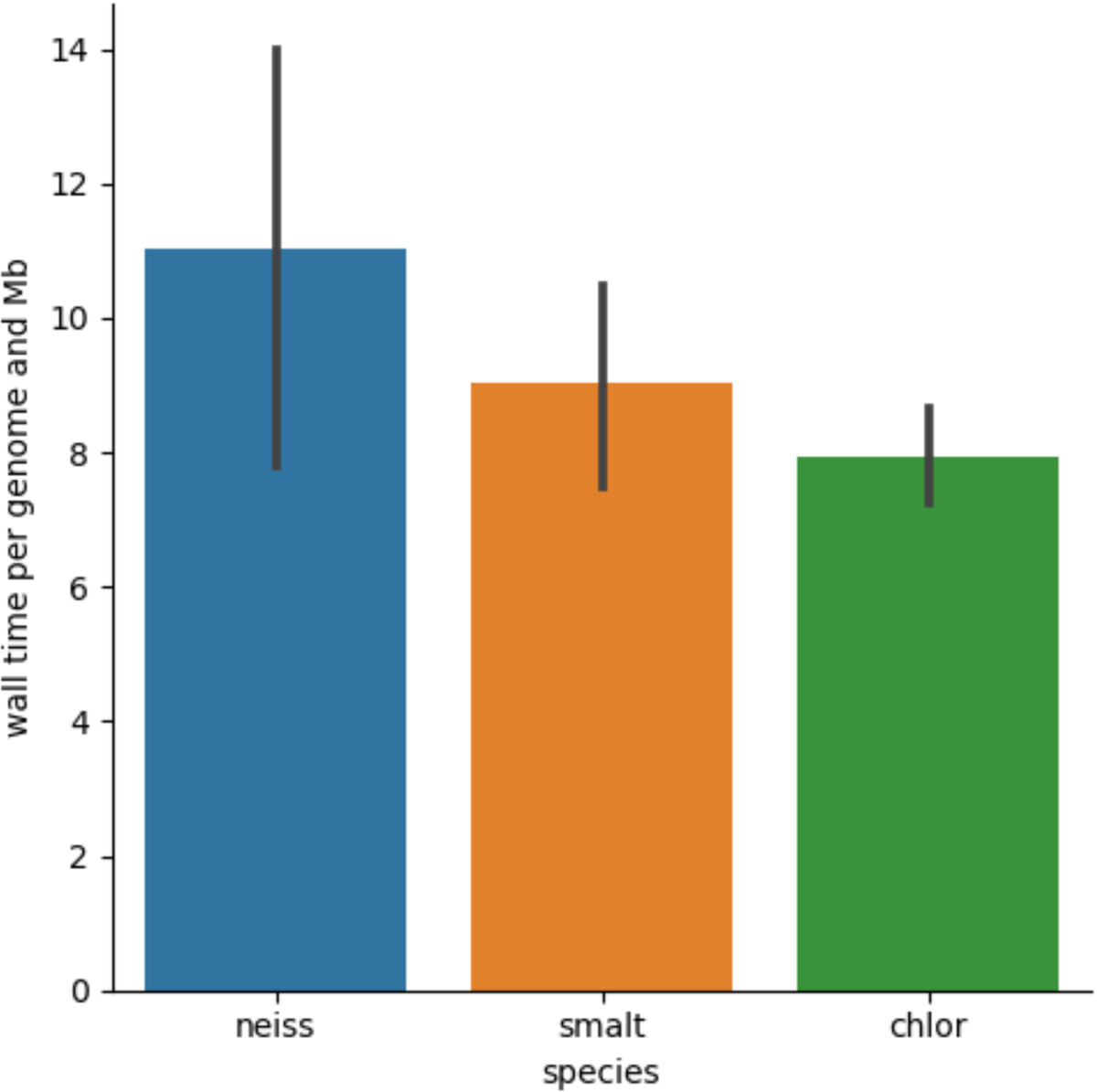
Approximate elapsed run time per megabase sequence length calculated from the same runtime data generated in **Supplementary Figure 3**.

## Notes

http://rarefan.evolbio.mpg.de/

### Competing Interest Statement

The authors have declared no competing interest.

### Summary of Updates

Added PCI Genomics badge.

